# Behavioral Thermoregulation in Captive Fish: Molecular, Physiological, and Welfare Implications

**DOI:** 10.1101/2023.10.10.561692

**Authors:** Nataly Sanhueza, Ricardo Fuentes, Andrea Aguilar, Beatriz Carnicero, Humberto Mattos, Yuniel Rubalcaba, Victoria Melin, David Contreras, Sebastian Boltana

**Author notes:** Correspondence: Sebastián Boltaña.

## Abstract

Environmental Enrichment (EE) serves as a cornerstone in the attempt to emulate natural habitats for captive organisms. While substantial strides have been made in this field, current methodologies still grapple with discrepancies between recreated habitats and the innate conditions vital for maintaining biological homeostasis in captive species. Our study highlights the pivotal role of behavioral thermoregulation in modulating molecular and physiological outcomes in captive fish. Collective evidence suggests that enabling fish to autonomously regulate temperature confers numerous beneficial cellular and systemic effects. Specifically, introducing a thermal gradient within the EE paradigm correlated with increased survival metrics, enhanced physiological parameters, and improved welfare indices, establishing the criticality of thermoregulation in captivity. In contrast, the lack of a thermoregulatory framework resulted in the emergence of transient free radicals, a clear marker of temperature-induced oxidative stress. Persistent disruptions in free radical equilibrium, especially in uniform temperature settings, were linked to DNA damage, heightened cellular apoptosis, tissue anomalies, and metabolic deviations. In conclusion, this research underscores the significance of behavioral thermoregulation as an integral feature of EE, especially related to fish in controlled environments. Our data present key biomarkers valuable for optimizing fish welfare and highlight the necessity for sustained research into their adaptability and survival benchmarks. Such insights aim to enhance EE protocols, fortifying their efficacy in mirroring natural habitats and, in turn, advancing the welfare benchmarks of captive organisms.

## Introduction

Environmental Enrichment (EE) is a pivotal strategy employed to enhance the welfare of aquatic organisms in captivity. Generally, EE primarily focused on augmenting the physical complexities of habitats. However, current research reveals a broader approach, incorporating sensory, social, and nutritional stimuli (Giménez-Candela et al., 2020). The primary objectives of EE extend beyond survival to promoting optimal sensory-motor interactions and mitigating stressors to prevent anomalous behaviors (Fraser, 2008). For a comprehensive understanding, EE’s impact on animal welfare can be classified into three categories: 1) Functional: Focusing on physiological health and metabolic function optimization. 2) Affective States: Minimizing stress and alleviating negative experiences such as pain. 3) Natural Living: Providing conditions that emulate natural habitats, fostering intrinsic capabilities like thermoregulatory adjustments (Fraser, 2008; Jena, 2017). Diving deeper into the EE framework, it is essential to consider the simulation of natural environmental factors in captivity, such as water temperature fluctuations. Water temperature, in particular, exerts significant influence over fish thermoregulatory behavior, directing their metabolic pathways (Abram et al., 2017). Ectothermic organisms like fish maintain thermoregulation by actively seeking optimal thermal environments aligned with their physiological needs (Žagar et al., 2018). Providing varied thermal gradients in captivity is vital to stimulate these behaviors, enhancing their functional capacities and optimizing biological processes. However, the challenge remains to effectively replicate natural thermal gradients in controlled settings. Existing research highlights the numerous benefits of the thermoregulation in such settings, pointing to decreased cortisol levels, reduced aggressive behaviors, and improved overall welfare metrics (Sanhueza et al., 2018).

In the context of animal welfare field, Environmental Enrichment (EE) stands as a cornerstone, profoundly affecting the well-being of captive species (Mellor et al., 2020). Enhanced welfare standards not only promise improved production outcomes but can also mitigate challenges such as antimicrobial interventions in aquaculture management. Such efforts not only provide to the immediate health of fish stocks but also combat challenges of antimicrobial resistance, with potential downstream benefits for human health (EMA & EFSA, 2017). Historically, welfare assessments in intensive aquaculture systems were largely short sighted, focusing mainly on immediate production indicators. Traditional approaches prioritized metrics like growth rates, somatic indices, and anatomical observations (Ellis et al., 2012). However, a more nuanced understanding has emerged, suggesting that welfare and productivity share a complex relationship over an organism’s lifespan. This ‘long-term welfare’ perspective considers the organism’s entire life cycle, from early development to maturity (Directive 2010/63/EU, 2009; FAWC, 2009). Given this shift, our study aims to develop, validate, and incorporate holistic welfare metrics, capturing the broader life experiences of captive fish. Moving beyond mere reactive indicators, such as mortality rates or visible anomalies, our goal is to establish a comprehensive and proactive set of benchmarks, advancing the field of aquatic welfare assessment.

Stress from captivity presents significant challenges for fish, often resulting in accelerated aging markers, such as DNA damage and telomere erosion. A core contributor to these outcomes is inadequate welfare conditions, leading to cellular deterioration and subsequent effects on health, reproduction, and survival rates (Fuller et al., 2021). A primary mechanism behind these detrimental effects is oxidative stress, arising from an imbalance between reactive oxygen species (ROS) production and the organism’s antioxidative responses. When ROS and free radical generation outpaces the antioxidative capacity, exemplified by enzymes like superoxide dismutase 1 (sod1) and glutathione peroxidase 1 (gpx1), cellular damage follows (Lushchak, 2011; Sies, 1985).

Environmental stressors enhance this situation. For example, pollutants, thermal fluctuations, and low oxygen levels can heighten ROS production, intensifying DNA impacting on the telomere damage (Blier, 2014; Harman, 1972). Specifically, aquatic pollutants can elevate 8-hydroxy-2’-deoxyguanosine (8-OHdG) levels, a DNA damage marker, as observed in species like the rainbow trout and golden grey mullet following heavy metal exposure (Alak et al., 2017; Oliveira et al., 2010). Additionally, contaminants such as cadmium can induce global DNA methylation shifts, potentially altering epigenetic patterns. This is evidenced by methylation changes in species like the European eel and rainbow trout, associated with variations in DNA methyltransferase mRNA levels (Reiser et al., 2021). Furthermore, specific stressors like elevated temperatures influence telomere lengths, as documented in the Siberian sturgeon and European chub (Simide et al., 2016; Molbert et al., 2021). Studies on zebrafish further confirm the susceptibility of telomeres to various stressors (Evans et al., 2021).

Finally, this study focuses on the influence of EE, highlighting thermoregulation, on long-term welfare metrics. In a thermoregulatory environment, our findings point to reduced oxidative stress and DNA damage in *Salmo salar*, alongside a notable positive correlation between cumulative mortality and global DNA damage and methylation. These findings present novel indicators for comprehensive welfare assessments in fish.

## 1. Materials and Methods

### 1.1. Fish Maintenance and Experimental Thermoregulation Conditions

All experimental protocols were conducted at the ThermoFish Lab, Biotechnology Centre, University of Concepcion, Concepcion, Chile, adhering to the research guidelines for laboratory animal use set by the Chilean National Commission for Scientific and Technological Research (CONICYT) and approved by the University of Concepcion Institutional Animal Care. Figure 1 provides a schematic representation of the thermal treatments, sampling strategies, and experimental procedures employed. Two advanced freshwater recirculation systems, equipped with UV sterilization and a flow rate of 5 m3/h each, were utilized. The fish received a commercial diet (INICIO PLUS followed by PLUS 18%, Biomar, SA, Puerto Montt, Chile) twice daily and were adjusted to a 12:12 h light-dark photoperiod (L:D) at a stable 12 ± 0.8 °C. Following a ten-day acclimation period, fish were randomized into two distinct thermal conditions, each with triplicate sets: A non-thermoregulatory group, maintained at a consistent temperature of 12.0 ± 0.7 °C. A thermoregulation group, exposed to a varied thermal gradient ranging from Tmin = 9.60 °C to Tmax = 16.4 °C, with a difference (ΔT) of 6.80 °C. The chosen temperature parameters were based on *S. salar* natural thermal range (7-22 °C as per Jobling (1981) and Torrissen et al. (2011)), and the constant temperature was determined in line with standard land-based aquaculture practices (Boltaña et al., 2017). Both groups were subjected to these conditions for a five-month duration, managed via an external water jacket system programmed at the specified temperatures.

**Figure 1.**
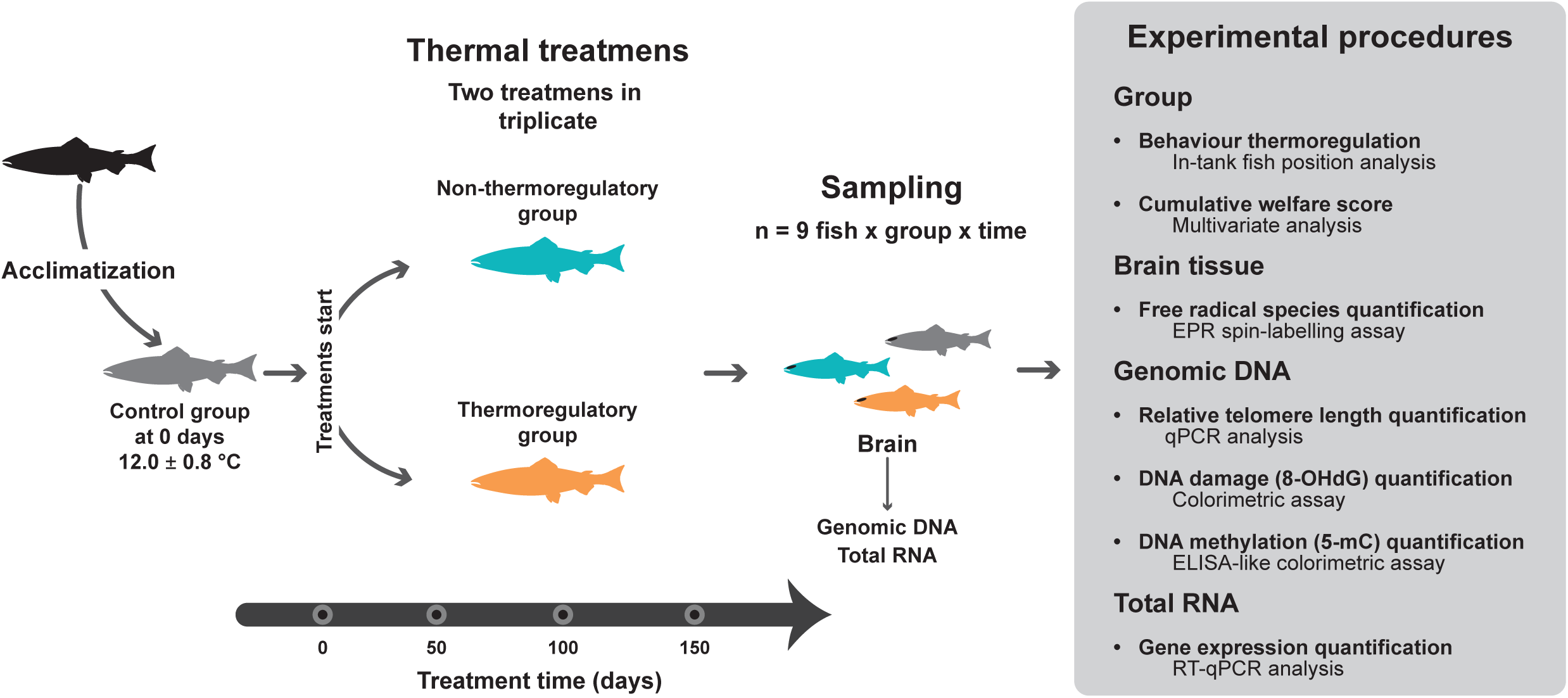
Experimental design, sampling, and subsequent procedures. The schematic shows the experimental design of thermal setup treatments for the rearing of *S. salar* for 150 days. The no-thermoregulatory group was kept at a constant temperature (12.0 ± 0.7 °C, cyan), and the thermoregulatory group was kept at a thermal gradient (T_min_ = 9.60 °C to T_max_ = 16.4 °C; ΔT 6.80 °C, orange). As a control group, 30 fish were sampled 0 days after starting treatment (dpt). At 50, 100, and 150 dpt, nine fish per thermal treatment and time were sampled. The diagram summarizes the analyzes performed to study the behavior thermoregulation and long-term welfare state (cumulative welfare) of experimental fish. EPR: electronic paramagnetic resonance; 8-OHdG: 8-hydroxy-2-deoxyguanosine (oxidized guanosine residues); 5-mC: 5-methyl cytosine (global DNA methylation).

### 2.2. Behaviour Analysis

The study on thermoregulatory behaviour involved tracking fish distribution within the distinct experimental setups: thermal gradient and constant temperature (as depicted in Fig. 2). Data collection occurred at specified intervals post-treatment: 0, 50, 100, and 150 days. During these intervals, temperature recordings were taken every 15 minutes, lasting 10 seconds each, accumulating to 96 readings daily. To monitor fish distribution, video surveillance was employed for a 24-hour duration. Image captures were taken every 15 minutes, yielding a total of 96 images daily. These images were subsequently processed using Image-Pro Plus 7 software (Media Cybernetics, Inc., Rockville, MD, United States) to determine the XY coordinates corresponding to the fish’s centre of mass. This data was then utilized to deduce individual thermal preferences, correlating fish positions with the respective temperatures.

**Figure 2.**
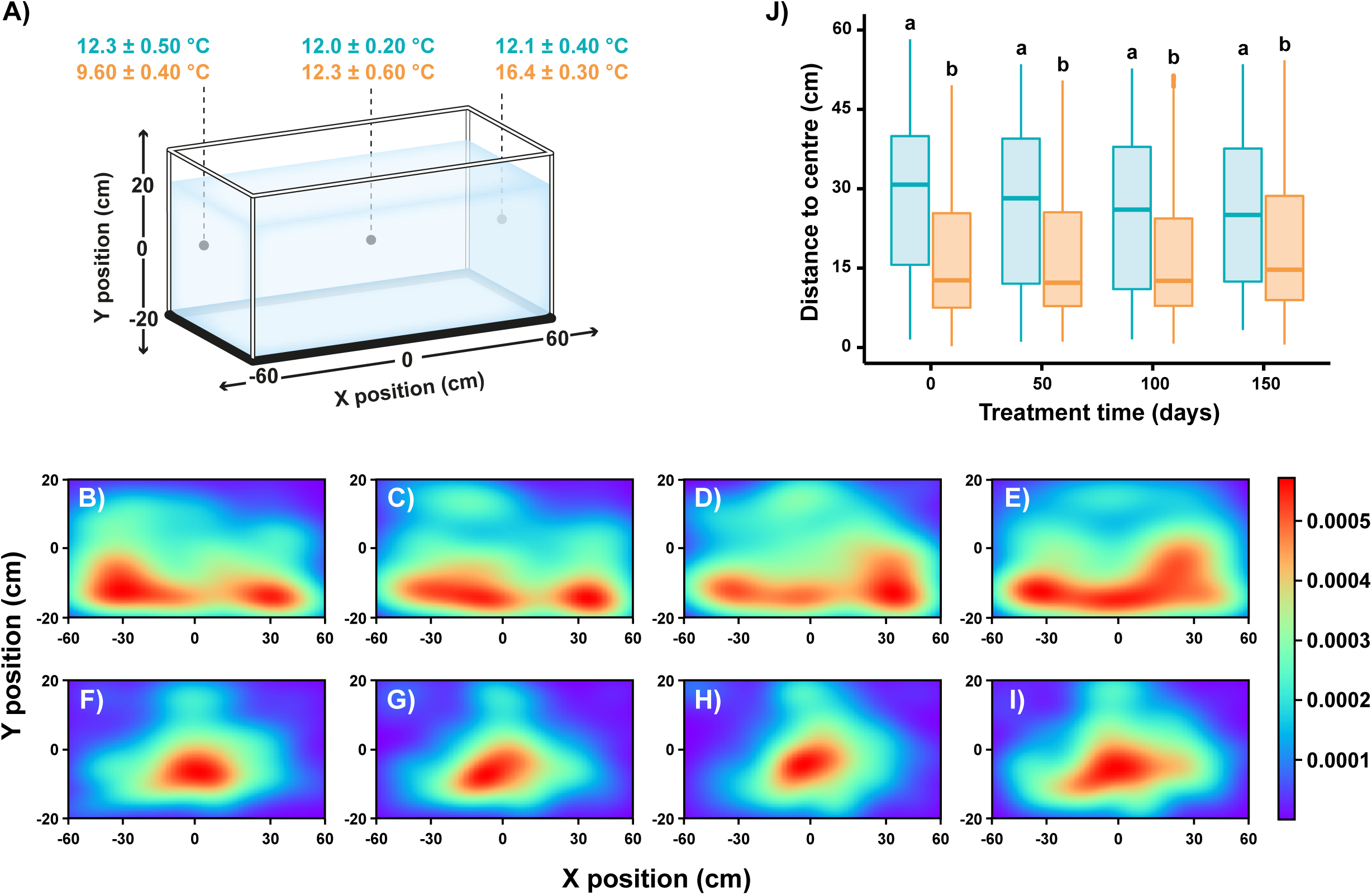
Thermoregulation behaviour of *S. salar* under two thermal treatments, constant temperature (no-thermoregulatory group, cyan) and thermal gradient (thermoregulatory group, orange). **(A)** The diagram shows the spatial (cartesian coordinates X, Y in cm) set-up of temperatures in the treatment tank. The centre of the tank was established at 0,0 coordinates. **(B and I) Color** maps representing the probability density function (Kernel density estimation) of fish position X, Y inside the constant temperature **(B and E)** and thermal gradient **(F and I)** tanks at 0 **(B and F),** 50 **(C and G),** 100 **(D and H)** and 150 dpt **(E-I)**. **(J)** Boxplot showing the position of the fish in the tank as a quantitative measure of individual thermal preference. The results are presented as the individual distance from the center of the tank at 0, 50, 100, and 150 dpt. Different letters denote significant differences between groups (mean ± SD, Kruskal-Wallis test followed by a Wilcoxon rank sum test, p > 0.05).

### 2.3. Free radical species measurement by electronic paramagnetic resonance analysis (EPR) spin trapping

The analysis of free radical species was carried out using an electronic paramagnetic resonance (EPR) EMX micro 6/1 Bruker spectrometer. EPR equipment worked in band X equipped with a Bruker Super High QE resonator cavity and 5,5-dimethyl-1-pyrroline N-oxide (DMPO) as a spin trap. For this analysis, brain tissue (20 mg) was homogenized in 100 µl 1X PBS (Gibco, Thermo Fisher Scientific, Waltham, MA, United States). The homogenized samples were then mixed with a 100 mmol / l DMPO solution in dimethylsulfoxide (DMSO). Measurements were made at 20 ° C in an EPR cell for solids using 50 µl of sample/solution in a sealed capillary. Instrumental conditions were the following: field centre 3515 G; sweep width 80 G; microwave power 20 dB; frequency modulation 100 kHz; constant time 0.01 ms; weep time 10 s; amplitude modulation, 1.00 G; and receiver gain 30 dB. 30 spectral accumulations were made for each sample. Simulation and adjustment of the EPR spectra were performed using an online simulator (https://www.eprsimulator.org/isotropic.html).

### 2.4. Genomic DNA and Total RNA Isolation

Genomic DNA (gDNA) was extracted from brain samples (25 mg) with the DNeasy blood and tissue kit (Qiagen, Hilden, Germany) according to the manufacturer’s protocol and resuspended in 20 µl of DEPC water. We evaluated the quantity and quality of DNA with a spectrophotometer/fluorometer (DeNovix DS-11; DeNovix Inc. Wilmington, USA) and the integrity was confirmed by denaturing agarose gel electrophoresis. Only samples with an A260/280 ratio between 1.8 and 2.1, and an A260/230 ratio above 1.8 were used to prepare a working stock of 50 ng/μl in DEPC water. Total RNA was extracted from brain samples (100 mg) with TRIzol reagent (1 ml; Invitrogen, Carlsbad, CA, USA) according to the manufacturer’s indications and resuspended in 20 µl of DEPC water. RNA samples were quantified by absorbance at 260 nm and integrity was confirmed by denaturing agarose gel electrophoresis. Only samples with an A260/280 ratio between 1.8 and 2.1, and an A260/230 ratio above 1.8 were used to prepare a working stock of 200 ng/μl in DEPC water.

### 2.5. Relative quantification of mRNA

cDNA was synthesized from total RNA (200 ng/µl) extracted from brain tissue using the reverse transcription supermix iScript TM for RT-qPCR (Bio-Rad, Carolina, United States) according to the manufacturer’s indications. RT-qPCR was performed using the Bio-Rad CFX 96 real-time system. qPCR analysis was performed using 1x SsoAdvanced Universal SYBR Green Supermix (Bio-Rad). The cDNA used in the qPCR assays was first diluted with nuclease-free water (Qiagen, Hilden, Germany). Each qPCR mixture contained SsoAdvance Universal SYBR^®^ Green Supermix, 2 µl cDNA, 500 nM of each primer, and RNase-free water to a final volume of 10 µl. A real-time PCR assay was performed to analyse the expression pattern of *superoxide dismutase 1 (sod1)* and *glutathione peroxidase 1 (gpx1)* in salmon brain tissue. The comparative ΔΔCt relative expression of Ct relative to Ct and the *elongation factor-α* (*ef-α*) as a housekeeping gene were used. The details of the primers are given in the supporting information (Table S1).

### 2.6. Oxidative DNA damage (8-OHdG) quantification assay

Oxidative DNA damage quantification was carried out using the EpiQuikTM 8-OHdG DNA damage quantification direct kit (Epigentek Group, Inc., Farmingdale, NY USA). This is based on colorimetric quantification of oxidized guanosine residues (8-hydroxy-2-deoxyguanosine; 8-OHdG), a sensitive marker of oxidative DNA damage and oxidative stress (Kasai, 1997; Pilger & Rüdiger, 2006). The procedure was carried out according to the manufacturer’s instructions. The absorbance (OD) was read at 450 nm using a microplate reader (BioTek™ 800™ TS; BioTek Instruments, Inc., Winooski, VT, USA). The samples were calibrated with the 8OHdG standard at 5 pg/µl to 200 pg/ µl. The standard characteristics of the curve of the curve were slope = 7.801, y intercept = 0.052 and R^2^ = 0.992, and the coefficient of variation between the tests between the tests between the tests (samples in duplicate) was 2.13%. Oxidative DNA damage levels were calculated as the percentage of genomic DNA containing 8LOHdG (8LOHdG % = absolute amount of 8OHdG / total amount of DNA × 100).

### 2.7. Global DNA methylation quantification assay

DNA methylation levels were measured using the MethylFlash TM Global DNA Methylation (5-mC) ELISA Easy Kit (Epigentek) following the manufacturer’s recommendations. Briefly, 100 ng of brain sample gDNA was used for the quantification of 5-methylcytosine (5-mC). Samples were read colorimetrically using a microplate reader (BioTek™ 800™ TS; BioTek Instruments) at 450 nm absorbance. To calculate the percentage of methylated DNA, a standard curve was first generated and the OD values versus the positive control (containing 5% 5mC at 5 ng/µl, 50 µg/ml) at each percentage point. The standard characteristics of the curve were slope = 0.134, y intercept = 0.388 and R^2^ = 0.980, and the coefficient of variation between the assays (samples in duplicate) was 3.06%. Finally, the percentage of methylated DNA (5 mC) in total DNA was calculated using the formula of the reported manufacturer.

### 2.8. Relative telomere length assay (RTL)

Relative telomere length (RTL) was measured using quantitative real-time PCR (qPCR) (Cawthon, 2002), and applicable MIQE guidelines (Bustin et al., 2009) were followed to ensure a high level of quality and reproducibility in qPCR analyses. The RTL was determined as the number of repeats of telomeres (T) per number of copies of the reference gene (S). Telomere repeats and *glyceraldehyde 3-phosphate dehydrogenase* (*gap8*) as a reference gene were amplified from brain genomic DNA as published (Cawthon (2002); McLennan et al. (2016), see Table S1). The DNA of the reference sample was a pool of all samples. Standard curves were generated using the CFX Maestro software v.1.1 (Bio-Rad) and the PCR efficiency (E) calculated as E = 10^[–1/slope]^. The standard curve characteristics (slope, y intercept and R^2^) and E of all plates are presented in Table SII. The relative length of the telomere length (T/S ratio) was calculated using the mathematical model for relative quantification by Pfaffl (2001; Pauliny et al., 2015):

### 2.9. Measurement of Long-Term Welfare

Long-term welfare was examined with a multivariate analysis according to (Turnbull et al., 2005). In summary, principal component analysis was used to combine four cumulative welfare metrics into a single score. PCA was performed using the prcomp function from the Stats R package. Oxidative DNA damage (8-OHdG %), DNA methylation (5-mC %), relative length of the telomere (T / S ratio), and cumulative mortality (%) at 50, 100 and 150 dpt were used as dependent variables. Factor scores from the first principal component as a single and integrated value were used, which increase with increasing cumulative welfare. PCA visualization was carried out using the fviz_pca_biplot function from the factoextra R package.

### 2.10. Statistical analysis

All statistical evaluations were performed using RStudio software (R Development Core Team, version 4.0.3). Before diving into in-depth statistical methods, the data underwent normality and variance homogeneity checks using the Shapiro-Wilk and Levene tests, respectively. For understanding thermoregulatory behaviour, we employed kernel density estimation (KDE) to establish the probability density function, drawing from the fish’s spatial positioning (XY coordinates) inside the tanks. Alongside, the average distance each fish maintained from the tank’s midpoint was calculated. These distances were then evaluated using a Kruskal-Wallis test, followed by a Wilcoxon rank sum test for detailed post-hoc analysis. Multiple biometric indicators, such as radical/DMPO relative concentration, gene expression ratios, 8OHdG levels, T/S ratios, and 5-mC percentages, were analyzed using two-way ANOVA. The variables considered were the type of treatment and time (expressed as days after the treatment commenced). Welfare evaluations followed a similar pattern, with a two-way ANOVA and subsequent inspection using Tukey’s HSD tests. To explore the relationships between cumulative mortality and metrics like 8OHdG concentrations, 5-mC percentages, and T/S ratios, we applied linear regression. This was supported by calculating Spearman’s correlation coefficient (rs). For visualizing our findings, we utilized the ggplot2 package in R and further refined visual data using GraphPad PRISM v6.0 (GraphPad Software, Inc., California, USA).

## 2. Results

### 2.1. Spatial Behavior of *S. salar* in Different Thermal Environments

To understand the thermoregulatory behaviors of *S. salar*, we studied their positioning preferences in two distinct environments: a dynamic thermal gradient and a uniform temperature setting (Fig. 2A). Monitoring their movements at 0, 50, 100, and 150 days revealed notable differences in behaviors between the two groups. In the consistent temperature environment, fish predominantly occupied the tank’s bottom section (Y position, 5-20 cm), demonstrating a wide range of lateral movements (X position from -20 to 20 cm) as shown in Fig. 2B-E. However, in the dynamic thermal gradient, fish displayed a preference for the tank’s central region, mostly within the Y position > -15 cm and an X range of -30 to 30 cm. This behavior likely reflects their tendency to stay within an optimal temperature zone (Fig. 2F-I). To account for potential influences of tank design and features on the fish’s vertical distribution, we computed the average positional deviation from the tank’s geometric center (XY 0.0) (Fig. 2J). Analysis using Kruskal-Wallis (p = 2.2 × 10^-16) and Wilcoxon range tests highlighted marked positional variations in the non-thermoregulatory group, suggesting a wider movement range (9, 6 to 16.4 °C, position X spanning from -60 to 60 cm). This expansive movement is likely attributed to the unrestricted thermal setting. Conversely, fish in the thermoregulatory environment remained largely centralized, possibly avoiding extreme temperatures near the tank’s peripheries.

### 2.2. Oxidative Stress Responses in S. salar Across Thermal Environments

Oxidative stress results from an imbalance in the production and detoxification of reactive oxygen species (ROS) within cellular systems. Environmental factors, such as temperature variations, can disturb this balance, thereby impacting biological homeostasis and ROS production. To gain insights into how *S. salar* responds to these changes, we assessed oxidative stress markers in the fish brain using two methodologies. First, we utilized Electron Paramagnetic Resonance (EPR) to quantify the generation of free radicals (Fig. 3). Our EPR data, highlighted in Figs. 3A-D, indicated comparable paramagnetic resonance spectra across both thermal settings. However, a gradual decline in signal intensity was noted over time. Further, a two-way ANOVA of the spectral curve’s area demonstrated a significant reduction in radical content from 50 to 150 days post-treatment (dpt) in both thermal environments (Fig. 3E). Secondly, to understand the antioxidant defense mechanisms in place, we examined the expression levels of antioxidant enzymes, *sod1* and *gpx1*, using RT-qPCR (Fig. 4). Our findings revealed an up-regulation of these enzymes in fish exposed to the thermoregulatory environment. Specifically, the *sod1* transcript levels at 100 and 150 dpt were markedly elevated in fish from the thermal gradient compared to those at a steady temperature (Two-way ANOVA treatment * time, p = 1.52 × 10^-2, Fig. 4A). In contrast, the constant temperature group displayed stable *sod1* transcript levels throughout the study duration (Fig. 4A). The *gpx1* mRNA levels also showcased distinct patterns between the two groups, with the thermoregulatory environment leading to increased transcript levels over time, especially at 50 and 150 dpt, compared to the constant temperature setting (two-way ANOVA p = 4.08 × 10^-2, Fig. 4B).

**Figure 3.**
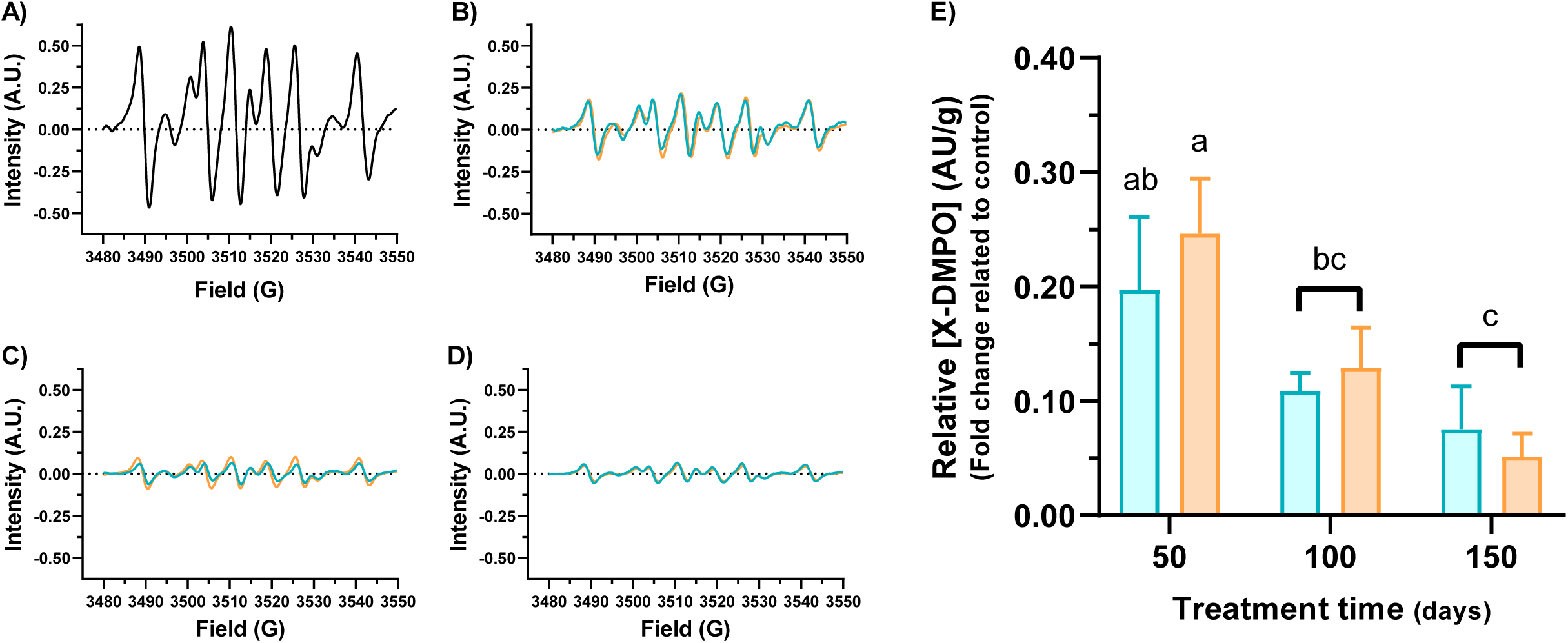
Free radicals generated by oxidative processes in the brain of *S. salar* under two thermal treatments, constant temperature (no-thermoregulatory group; cyan line or bars) and thermal gradient (thermoregulatory group; orange lines or bars). **(A-D)** Electronic paramagnetic resonance spectra of DMPO spin adducts at 0 (control untreated), 50, 100, and 150 dpt, respectively. **(E)** Bar graph showing the relative concentration of DMPO spin adducts. The results are presented as the fold change in the relative concentration of radical/DMPO (arbitrary units, AU) per tissue sample mass ([X*DMPO] AU/g) at 50, 100 and 150 dpt over untreated control. Different letters denote significant differences between groups (mean ± SD, two-way ANOVA, p > 0.05).

**Figure 4.**
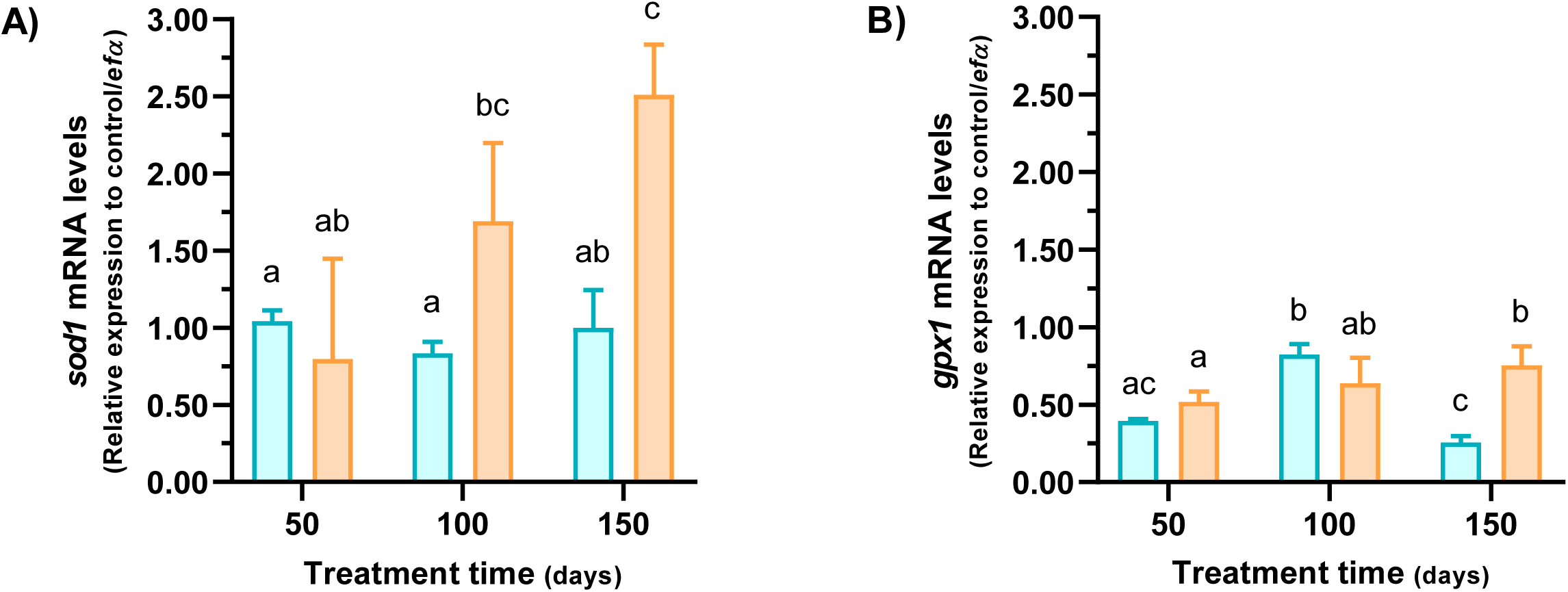
Expression profile of antioxidant enzymes in the brain of *S. salar* under two thermal treatments, constant temperature (no-thermoregulatory group; cyan bars) and thermal gradient (thermoregulatory group; orange bars). Bar graph showing the differences in *sod1* **(A)** and *gpx1* **(B)** mRNA levels between treatments at 50, 100, and 150 dpt. The results are presented as relative expression to the untreated control (0 dpt) and *ef-α* as the housekeeping gene (mean ± SD, two-way ANOVA following Tukey’s post hoc test, p < 0.005). Different letters denote significant differences between groups.

### 2.3. Influence of Thermoregulatory Environments on DNA Structural Integrity

Oxidative stress, precipitated by the reactive oxygen species (ROS), results in cellular damage and perturbations in tissue function. These ROS, characterized by their elevated reactivity, are implicated in oxidative deleterious effects on DNA, lipids, and proteins (Harman, 1956). To elucidate the ramifications of thermal conditions on oxidative balance, we develop on an analytical exploration of DNA damage within the cerebrum of the fish. Our primary investigational approach was to discern the interrelationship between DNA damage and oxidative stress within fish and the thermoregulatory comportment. Employing the metric of 8-OHdG to quantify oxidized guanosine residues, a significant differential emerged between fish subjected to thermoregulatory conditions versus those in a consistent temperature environment (two-way ANOVA, p = 9.66 × 10^-3). As depicted in Fig. 5A, fish in homogenous temperature conditions exhibited an augmented extent of DNA damage, especially pronounced at 50 and 150 dpt, when juxtaposed against their thermoregulatory counterparts.

**Figure 5.**
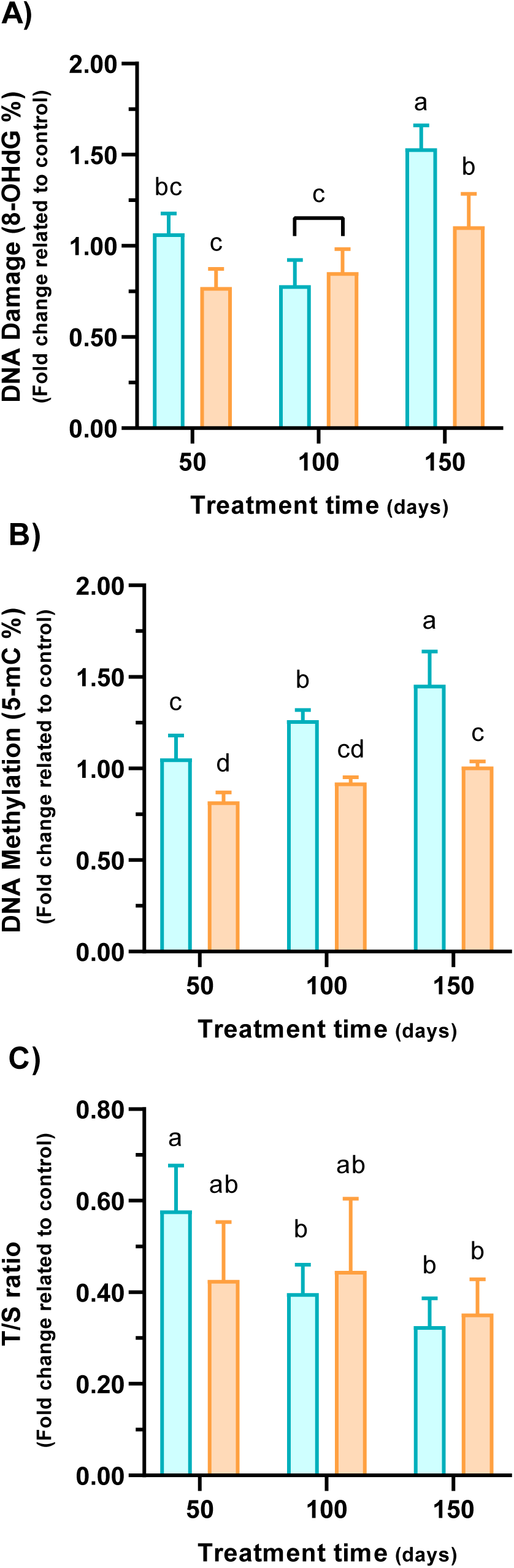
Aging-related hallmark levels in the brain of *S. salar* under two thermal treatments, constant temperature (no-thermoregulatory group; cyan bars) and thermal gradient (thermoregulatory group; orange bars). The bar graphs show differences in oxidative DNA damage as the percentage of 8-OHdG **(A)**, methylated DNA as the percentage of 5-mC in total DNA **(B),** and relative telomere length as the T / S ratio **(C)**. The results are presented as the fold change in 8OHdG%, 5-mC%, or T/S ratio at 50, 100, and 150 dpt over the untreated control at 0 dpt (mean ± SD, bidirectional ANOVA following Tukey’s post hoc test, p = 9.66 × 10^-3^, p = 6.20 × 10^-3^ and p = 0.96, respectively). Different letters denote significant differences between groups.

Follow, we looking for on the DNA methylation, a pivotal mechanism in early somatic cellular differentiation and related with DNA repair processes (Sen et al., 2010). Extracted DNA from fish cerebral samples across both thermal conditions revealed distinct methylation profiles (two-way ANOVA, p = 2.05 × 10^-4, Fig. 5B). Notably, fish maintained in unvarying temperature conditions consistently manifested elevated 5-mC levels at 50, 100, and 150 dpt, in stark contrast to their counterparts in the thermal gradient. This data accentuates the potential of thermoregulatory behavior to induce a hypomethylative state within fish DNA (Fig. 5B). Nevertheless, the impact on transcriptional processes necessitates more comprehensive epigenetic assessments. Additionally, the relative telomere length, denoted as the T/S ratio, was ascertained for fish across both environmental conditions (Fig. 5C). It was observed that fish within a uniform temperature setting experienced a sequenced diminution in telomere length, with conspicuous disparities discernible between 100 and 150 dpt (two-way ANOVA p = 7.97 × 10^-2). In stark contrast, fish within the thermoregulatory environment maintained a uniform telomere length across the designated temporal intervals (50, 100, and 150 dpt). This data suggests that while homogenous temperature conditions may promote telomere erosion, thermoregulatory environments potentially offer a safeguarding influence on telomere preservation.

### 3.4 Thermoregulatory Environments and Implications for Fish Welfare

Building on our findings of temperature-induced DNA damage, we next investigated its implications on fish welfare. We utilized Principal Component Analysis (PCA) to consolidate data related to DNA damage, methylation, telomere shortening, and survival rates into composite welfare scores. As depicted in Figure 6A, the first two principal components accounted for 83.19% of the overall variance—62.44% and 20.76%, respectively. The first component was significantly influenced by metrics like % 8-OHdG (-0.51), % 5mC (-0.58), and % cumulative mortality (-0.54), whereas the telomere T/S ratio length (0.32) exhibited a positive loading. Subsequent two-way ANOVA analyses showed that the PC1 scores, representing cumulative welfare scores, were affected by the thermoregulatory conditions, duration of exposure, and their interaction (p-values: 4.55 × 10^-6, 4.16 × 10^-7, and 4.65 × 10^-3, respectively). Fish in the thermoregulatory setting displayed better welfare scores compared to those in non-thermoregulatory conditions (Fig. 6B). Over time, both groups exhibited a decline in welfare scores, but the constant temperature group had a more pronounced reduction, especially at the 100 and 150 dpt times (Fig. 6B). In further analyses, we used linear regression with Spearman correlation to explore relationships between cumulative mortality, oxidative DNA damage (% 8-OHdG), DNA methylation (% 5-mC), and relative telomere length (T/S ratio). We found significant positive correlations between cumulative mortality and both % 8-OHdG (rs = 0.54, p = 0.02, Fig. 7A) and % 5-mC (rs = 0.85, p = 7.26 × 10^-6, Fig. 7B). However, the relationship between cumulative mortality and telomere length was negative, though not statistically significant (rs = -0.30, p = 0.23, Fig. 7C).

**Figure 6.**
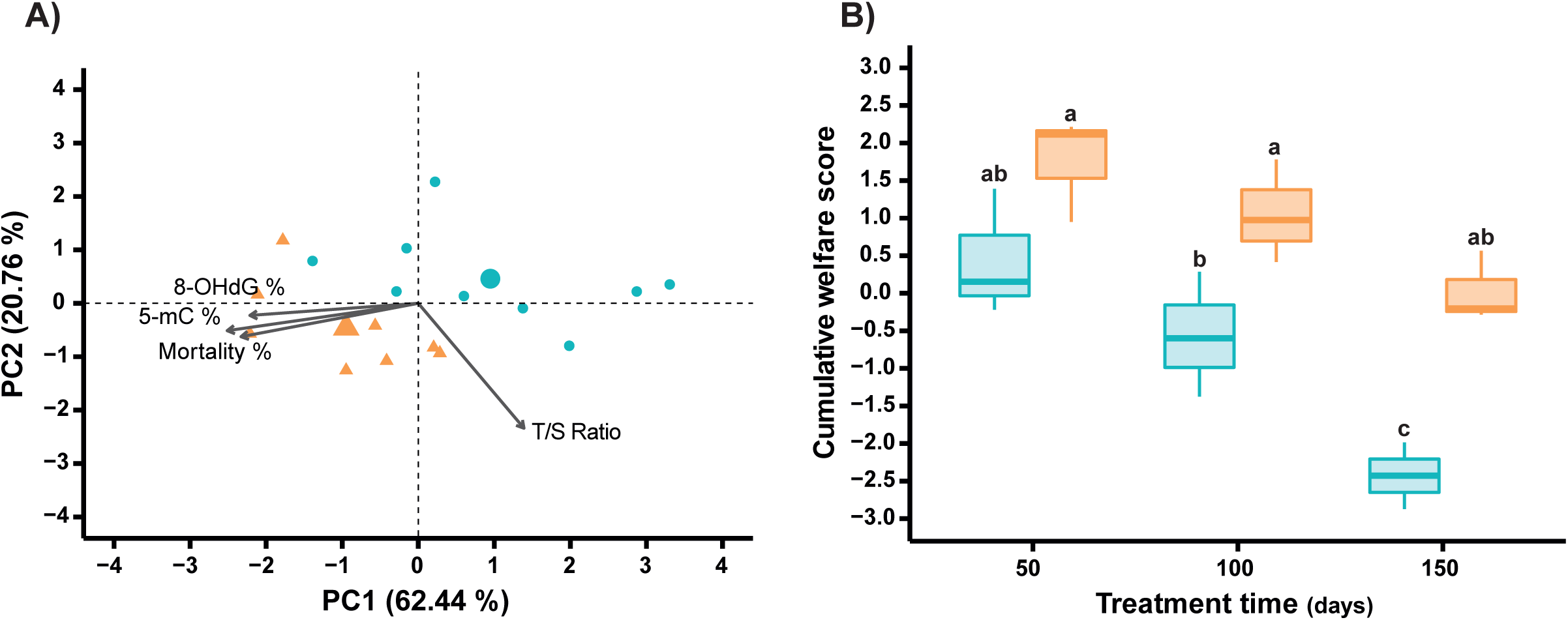
Long-term (cumulative) welfare state of *S. salar* under two thermal treatments, constant temperature (no-thermoregulatory group; cyan) and thermal gradient (thermoregulatory group; orange) at 50, 100 y 150 dpt. **(A)** Biplots generated from the analysis of principal components (PCA) for oxidative DNA damage (8-OHdG %), DNA methylation (5-mC %), relative telomere length (T / S ratio) and cumulative mortality (%). **(B)** Boxplot showing a cumulative welfare score that originated from factor scores for the first principal component. Different letters denote significant differences between groups (mean ± SD, two-way ANOVA following Tukey’s post hoc test, p = 4.65 × 10^-3^).

**Figure 7.**
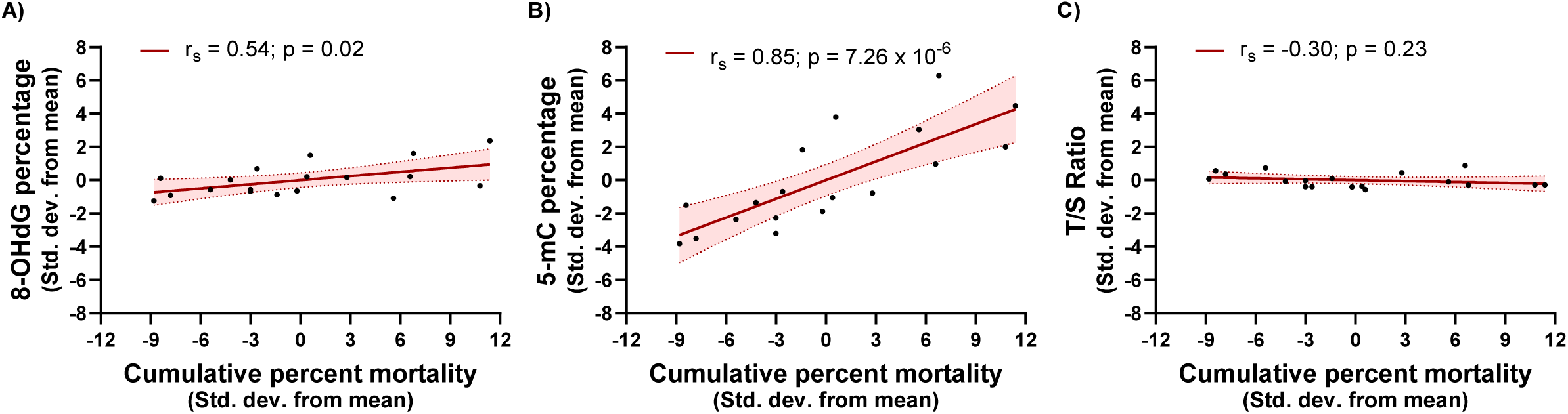
Relationship of cumulative mortality with the level of 8OHdG **(A), the percentage of** 5-mC (B), or the T / S ratio **(C)**. Results are presented as the standard deviation of the median values of each variable across thermal treatments, constant temperature (no-thermoregulatory group) and thermal gradient (thermoregulatory group) at 50, 100 y 150 dpt. A linear regression was used and the value of the Spearman correlation (rs) is shown on each graph.

## 3. Discussion

In captive systems or aquaculture conditions, physicochemical environmental conditions (e.g., water quality, temperature, photoperiod) are controlled within a relatively narrow range (Barreto et al., 2022). This control is prescribed in protocols to improve captive housing; however, it is essential to recognize which conditions can provide a significant source of EE and contribute to fish welfare. Here, we demonstrate that EE promotes positive welfare, focusing on the thermoregulatory and behavioural responses of fish in captivity. Interestingly, fish subjected to a thermoregulatory environment had lower mortality, oxidative stress, and DNA damage than those maintained at constant temperature. Our study highlights that temperature, as an EE tool, could influence the phenotypic traits and performance of Atlantic salmon in several ways. Compared to conventional fish reared at constant temperature, those grown under enriched conditions showed better survival and higher welfare parameters after 150 days of captivity. Since hatchery water comes from a common source, the observed differences between rearing methods are supposedly due to thermal treatment. Overall, these results agree with previous observations on salmonids (Boltaña et al., 2017; Sanhueza et al., 2018) and support the idea that a thermoregulatory environment drives the progression of better welfare conditions, and simple enrichment improvements can restrain this as the temperature settings in an aquaculture system.

An increase in oxidative stress is often linked when animals are under stressor conditions, such as fluctuations in environmental temperature (Nakano et al., 2014). On the basis of our EPR results, we found similar free radical production in both thermoregulatory groups, which implies a high availability of iron in the individuals. However, the regulation of free radicals of the fish group exposed to constant temperature was lower than that of the animals reared in the thermal gradient. Iron is essential for life; many proteins involved in crucial cellular processes require this mineral (Ajioka et al., 2006). An excess of intracellular iron catalyses the generation of free radicals through *Fenton chemistry* and causes damage to lipids, proteins, and DNA (Drakesmith & Prentice, 2008), leading to cell and tissue damage. Further exploration of genes associated with ROS reflects an increase in the antioxidant response in fish reared in the thermoregulation environment. We found an increase in the abundance of *sod1* and *gpx1*, which are closely related to the regulatory ROS response (Bayır & Bayır, 2017). This result highlights that the increase in oxidative stress as a result of constant temperature induces down-regulation of the mechanisms driving ROS homeostasis. ROS contributes to tissue damage in many pathophysiological conditions (Ryter & Choi, 2005). In vertebrates, an increase in ROS induces the activation of DNA damage (Czarny et al., 2018). Our findings also show that fish reared at constant temperature increase the percentage of 8-OHdG, which is a direct indicator of DNA damage (Pilger & Rüdiger, 2006). Several species of fish under stressor conditions also show dysregulation of antioxidative mechanisms (Tarifeño-Saldivia et al., 2018), however, further analysis is needed to dissect the regulatory network linking ROS and thermal-based dependence.

Interestingly, since free radicals and 8-OHdG levels are stressor indicators (Zhang et al., 2022), and both rates remained significantly higher in fish reared at constant temperature, confinement conditions would induce extensive cell damage. The non-thermoregulatory fish group showed the most significant DNA damage but within the normal range reported (Meier et al., 2020), which suggests that the increase in oxidative stress promotes controlled DNA damage as a mechanism to maintain the cellular metabolism (Martins et al., 2021). The lowest DNA damage observed in the thermoregulatory fish group might indicate that they reacted to temperature changes efficiently and responded appropriately to metabolic demands.

According to Huntingford et al. (2020), EE can transmit emotions and reveal perceptions and events from the external environment, improving biological functions. Thermoregulation stimulation can lead to physical, cellular, and social changes in individuals by activating brain regions responsible for motor movement execution (Tan & Knight, 2018). The unlike temperatures drive significant modifications in the metabolic performance of fish reared in a thermally changing environment over a long period (Aguilar et al., 2022). In this study, the persistence of the fish at a constant temperature led to profound modifications in the DNA methylation pattern. Increasing evidence shows that environmental challenges can affect the normal development of organisms by altering DNA methylation patterns (Berbel-Filho et al., 2019). Reprogramming of DNA methylation after fertilization is a dynamic mechanism essential for development and is considered sensitive to environmental stressors (Head, 2014). This epigenetic mechanism regulates transcription, ubiquitination, autophagy, and microRNA expression during early and late development (Hu, 2019), thus affecting several cellular behaviours (Rahman & McGowan, 2022). The present data show that non-thermoregulatory fish modify their global pattern of methylation. However, the DNA methylation assay used in this study only tested for a global change in the methylated cytosine content and did not detect changes at specific loci. Therefore, additional studies will determine which loci are affected by methylation processes under captive conditions.

Collective reports have revealed a close relationship between stressor conditions and telomere dynamics, showing that the effect of stressor situations results in substantial telomere attrition (Bateson, 2016; de Punder et al., 2019). Interestingly, numerous research groups have shown that adverse environmental conditions induce telomere attrition in mammals (Kotrschal et al., 2007; Stindl, 2004a), birds (Hall et al., 2004), reptiles (Olsson et al., 2018), and fish (Gopalakrishnan et al., 2013). However, these findings are inconsistent for all species, reflecting essential differences in species-specific biology. Specifically, our results showed that constant temperature induces attrition of the telomere. Assuming that the captivity conditions are similar in both treatments, the observed differences in telomere attrition are related to the thermal gradient response. Furthermore, the difference observed in the attrition of telomeres within species is not well understood, but has been associated with other stressor parameters such as the increase in oxidative stress, cortisol, hormones, or inflammatory molecules (Barrett & Richardson, 2011; Stindl, 2004b). Our results show a correlation between antioxidant response and telomere attrition (de Punder et al., 2019). This correlation is also observed in zebrafish exposed for 14 days to a stressful condition; where a lower cortisol level and antioxidant response show an impact on telomere length (Rambo et al., 2017). In our study, the stress factors detected in fish reared at constant temperature were higher than those kept in the thermoregulatory environment. In summary, our work indicates that individuals without access to EE develop poor welfare in response to stressful physiological conditions (Madaro et al., 2015).

## 4. Conclusions

The present data suggest that *salmon salar* under thermal environmental enrichment activates several regulatory responses to improve its welfare. The positive influence of thermoregulation on welfare parameters may be the adaptive behaviour of a protective response to stress. Furthermore, a positive correlation is found between the animal’s preference to explore new areas and the survival rate. Our current findings illustrate that the thermoregulation behaviour performed by the fish poses a temperature variation and choosing is a promising EE tool. This strategy allows organisms to avoid stress-induced effects in a captive environment. Increased levels of free radicals, DNA damage and methylation pattern, and telomere shortening in a non-thermoregulatory environment might be effective biomarkers to monitor animal welfare in laboratory and aquaculture housing.

## Author contribution statement

The study was conceived by SB. Behaviour experiments were performed by NS. NS, BC, AA, and HM performed a transcriptomic, ROS, and DNA damage analysis. DC and VM performed the EPR analysis. SB obtained the acquisition of funds. NS, RF, and SB drafted the manuscript with contributions from all other authors.

## Data availability statement

The original contributions presented in the study are included in the article/Supplementary Material.

